# PINK1 regulates cholesterol homeostasis via SCAP phosphorylation in human dopaminergic neurons

**DOI:** 10.1101/2025.08.27.672574

**Authors:** Karan Sharma, Claudia Cavarischia-Rega, Dina Ivaniuk, Christine Fröhlich, Jos F. Brouwers, Gerard J.M. Martens, Philip Seibler, Javier Jarazo, Thomas Gasser, Daniela M. Vogt Weisenhorn, Boris Macek, Julia C. Fitzgerald

## Abstract

Cholesterol is a key lipid enriched in neuronal membranes and essential for signaling and synaptic transmission. An imbalance in cholesterol levels may affect synaptic plasticity and contribute to neurodegeneration. Here, we identify in human dopaminergic neurons a mechanism linking loss of function of the Parkinson’s disease (PD) gene *PINK1* to altered cholesterol homeostasis. Loss of functional PINK1 impaired SCAP phosphorylation at Ser822 and Ser838, stabilizing SCAP and driving excess cholesterol biosynthesis. Cholesterol accumulated at the plasma membrane and in flotillin-rich lipid rafts, causing reduced neurotransmitter uptake and altering the distribution of dopamine transporter (DAT). Restoring PINK1 expression normalized cholesterol biosynthesis and levels. Moreover, the cholesterol-lowering drugs simvastatin and β-cyclodextrin rescued DAT distribution and neurotransmitter uptake defects. These findings demonstrate that PINK1 influences cholesterol homeostasis through SCAP phosphorylation at Ser822 and Ser838 and that restoring cholesterol levels mitigates phenotypes observed in PINK1 PD neurons. These findings further highlight the cross-talk between mitochondria and lipid homeostasis in PD models, underscoring the relevance of cholesterol levels to dopaminergic functions.

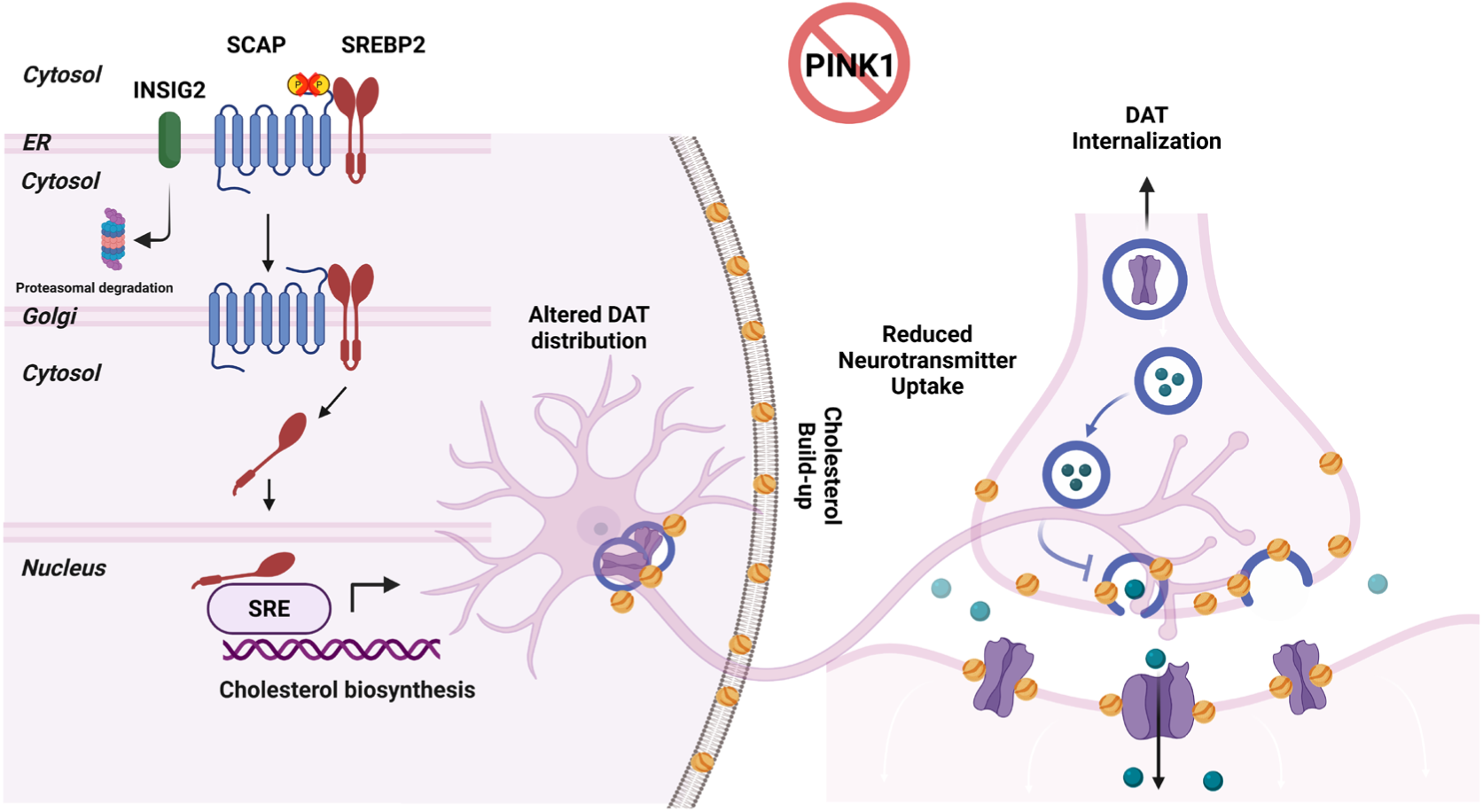
Graphical Abstract.

## Introduction

Mutations in phosphatase and tensin homolog-induced kinase 1 (*PINK1*) typically cause autosomal recessive Parkinson’s disease (PD)^1^. Clinically, PINK1-PD presents early (median age at 32 years) with tremor, bradykinesia and rigidity and responds well to levodopa treatment^2^. PINK1-associated neuropathology involves the selective degeneration of dopaminergic neurons (DaNs) in the *substantia nigra*, while Lewy body pathology is inconsistently observed^3^ ^4^.

The canonical function of PINK1 is mitochondrial quality control. Upon mitochondrial depolarization, PINK1 accumulates at the outer mitochondrial membrane, where it phosphorylates ubiquitin and Parkin, initiating an outside-in PINK1/Parkin-dependent mitophagy to remove damaged mitochondria^5–20^. Beyond mitochondria, PINK1 has been implicated in maintaining endoplasmic reticulum (ER) homeostasis by promoting selective clearance of the organelle^21^, regulating ER calcium release mediated by IP3R^22^ and modulating the ER unfolded protein response (UPR)^23^ ^24^. PINK1 loss of function (LOF) also disrupts ER-mitochondria contacts and calcium buffering, suggesting broader roles at ER-mitochondria interfaces ^25^.

Cholesterol, synthesized in the ER, is an essential structural and functional component of neuronal membranes. At the plasma membrane cholesterol supports neuronal signaling, vesicle trafficking and synaptic neurotransmission^26^. Cholesterol is synthesized *de novo* in the brain as cholesterol does not cross the blood brain barrier (BBB)^27^. Young and developing neurons make their own cholesterol, while adult neurons rely mainly on cholesterol imported from neighbouring astrocytes^28^ ^29^.

Cholesterol biosynthesis is tightly regulated by the sterol regulatory element-binding protein (SREBP) pathway, with sterol regulatory element-binding protein cleavage-activating protein (SCAP) as a key regulator^30^. During synthesis, insulin-induced gene (INSIG) proteins are dissociated from the SCAP-SREBP2 complex at the ER membrane^30–33^. The SREBP2-SCAP complex then leaves the ER and translocate to the Golgi, where SREBP2 is cleaved by site 1 and site 2 proteases (S1P and S2P, respectively). Cleaved SREBP2 then goes to the nucleus and binds to the sterol regulatory element (SRE) to transcribe SREBP2 target genes i.e., 3-hydroxy-3- methylglutaryl-CoA reductase (HMGCR) and squalene monooxygenase (SQLE) that are the rate-limiting enzymes for cholesterol biosynthesis^30^ ^34^. The activity of SCAP is negatively regulated by five sterol-sensing transmembrane (TM) helices (TM domains 2-6) and by phosphorylation involving the atypical protein kinase C λ/ι (PKCλ/ι) that phosphorylates multiple serine residues of SCAP at the C-terminus^35^.

Alteration of key proteins involved in cholesterol regulation, levels in blood and cellular distribution have been implicated in neurodegenerative diseases. In PD, the role of cholesterol in disease progression is unclear as there are conflicting reports about the association between serum cholesterol levels and PD risk^36–41^. However, the levels of cholesterol and their cellular impact in PD neurons remain unexplored.

Here, we investigated cholesterol regulation in PINK1 LOF DaNs. An unbiased phosphoproteomics screen revealed reduced phosphorylation of SCAP at Ser822 and Ser838 in PINK1 knockout (KO) and PINK1-Q456X PD neurons. Reduced regulation by phosphorylation, stabilized SCAP, enhanced cholesterol biosynthesis, and led to cholesterol accumulation at plasma membrane and in Flotillin 1 (FLOT1) rich lipid rafts. Elevated cholesterol disrupted dopamine transporter (DAT) distribution and impaired neurotransmitter uptake. Notably, both genetic restoration of wildtype (WT) PINK1 and pharmacological cholesterol lowering with simvastatin or β-cyclodextrin (βCD) rescued these phenotypes. Our findings identify regulation of cholesterol homeostasis as a previously unrecognized role of PINK1, mediated through SCAP phosphorylation, and suggest that targeting cholesterol metabolism may improve neuronal function in PINK1-linked PD.

## Materials and methods

### Ethics statement

The study was approved by the ethics committee (Institutional Review Board) of the Medical Faculty of the University of Tübingen and the University Clinic Tübingen (146/2009BO1 and 102/2005). A general consent form of the Hertie Institute for Clinical Brain Research Neurobiobank and a project-specific consent form including an information sheet about the study were used. All participants gave written, informed consent. All research adheres to the current, updated version of the Helsinki declaration.

### Cell culture

#### DaN differentiation

PINK1 wildtype (WT) and knockout (KO) iPSCs were previously generated and characterized in Bus et al.^42^, PINK1 Q456X and gene corrected (GC) iPSCs from PD patients #1 and #2 were previously generated and characterized in Jarazo et al.^43^. The PINK1 Q126P patient line was previously described in Prestel et al.^44^ and gene corrected by our group. These iPSC lines were differentiated by chemical induction using small molecules based on the protocol from Reinhardt, et al,^45^ with slight modifications^42^. The experiments were performed on day 21 post-differentiation start.

#### HeLa cell culture

WT and PINK1 W437X HeLa cells, characterized in Wettengel et al.^46^ were maintained in Dulbecco’s Modified Eagles Medium: 4.5 g/L glucose (D6429; Sigma-Aldrich) and supplemented with 10% fetal bovine serum and 1% penicillin/streptomycin.

#### Generation of PINK1 Q126P Gene Corrected iPSCs

The homozygous PINK1 Q126P iPSCs were cultured in mTeSR1 Plus medium (Stem Cell Technologies) on Matrigel (Corning). Endogenous expression of Cas9 was established by introducing a doxycycline-inducible Cas9 construct (Addgene 12551,^47^) within safe-harbour AAVS1 locus using plasmid-mediated HDR and Crispr-Cas9 in homozygous PINK1 Q126P iPSCs. A detailed description is given in the Supplementary methods.

### PINK1 KO mice

PINK1 KO mice, previously described in Glasl et al. ^48^, were used at 4 months (young) and 16 months (old) of age and bred in accordance with the regulations from the government of Upper Bavaria. The mice were kept on a C57BL/6J background and on an inverse 12-h light/12-h dark cycle (lights off at 18:00). Mice were provided with ad libitum access to standard chow and water. Mice were killed by cervical dislocation and the brains immediately removed. Thereafter, single-brain regions were dissected and the tissue shock frozen in liquid nitrogen.

### Cell Treatments

#### Cycloheximide treatment

To check for protein stability, cells were treated with 100 μM cycloheximide solution (CHX; 239765; Calbiochem, Merck Chemicals GmbH, an affiliate of Merck KGaA, Darmstadt, Germany) with an equal volume of DMSO used as a vehicle.

#### Mitophagy induction

Mitochondria were depolarized using 100 μM Antimycin A (#A8674-50MG Merck). Equal volumes of 95% ethanol were added as vehicle.

#### Simvastatin

Simvastatin was used at 10 μM concentration. The powder (#S1792, Selleckchem) was resuspended in ethanol and stored at -80°C. For each experiment, 200 μL of this solution was added to 300 μL 0.1N NaOH solution and heated for 2h at 50 °C. Then the pH was adjusted to 7 with HCl and the total volume was adjusted to 1.431 mL. This final solution was freshly prepared before each experiment and kept in the media for 16 hours.

#### β-cyclodextrin (βCD)

A 1 mM concentration of (2-Hydroxypropyl)-β-cyclodextrin (#H107-5G, Sigma-Aldrich) was freshly made with culture media and added to the cells for 16 hours.

#### Cholesterol

100 μM cholesterol (#C3045, Sigma) freshly resuspended in culture media was used to treat the cells and kept for 16 hours.

#### DAT inhibitor

10 μM of DAT inhibitor (GBR 12935 dihydrochloride, #0514, Tocris) was resuspended in DMSO and added to cells for 16 hours.

### Measurement of Free Cholesterol

The Total Cholesterol Assay Kit (Fluorometric) (STA-390, Cell Biolabs Inc) was used to measure free cholesterol (non esterified) according to manufacturer’s instructions. Absolute cholesterol levels were measured using cholesterol standards and normalized to protein input in the lipid extraction. For the mouse brain tissues, free cholesterol levels were normalized to the tissue mass used.

### Mass-Spectrometry-Based Phosphoproteomics

For each phosphoproteomics experiment, 1 mg total protein per sample was used. Proteins were reduced with 1 mM dithiothreitol (DTT), which was incubated for one hour at room temperature (RT) while shaking. To stabilize the reduction, 5.5 mM iodoacetamide (IAA) was added and incubated for one hour at RT while shaking in the dark. Samples were pre-digested with Lysyl Endopeptidase (LysC, Wako Chemicals) for 3 hours at RT. Next, four volumes of water were added and proteins were digested with trypsin (Promega Corporation) overnight. The reaction was stopped the following morning by acidifying the samples using around 0.1% v/v of trifluoroacetic acid. The peptides for proteome and phosphoproteome analysis were desalted and purified using C18 StageTips (Empore). Samples were measured on an Exploris 480 mass spectrometer (Thermo Fisher Scientific) online-coupled to a VanquishNeo UHPLC (Thermo Fisher Scientific). Chromatographic separation was performed on a 20 cm long, 75 µm inner diameter analytical HPLC column (ID PicoTip fused silica emitter; New Objective), packed in-house with ReproSil-Pur C18-AQ 1.9-μm silica beads (Dr Maisch GmbH). The raw data were processed using the MaxQuant program (version 2.2.0.0). The raw spectra were searched against the UniProt Homo sapiens database (104556 entries, downloaded 30.01.2024). The downstream analysis of MaxQuant output data was performed in Perseus (version 2.0.10.0). Contaminants, reversed hits and proteins identified only by site were filtered out. Scatter plots were prepared to assess reproducibility between replicates and Pearson’s correlation was calculated. For the proteome, Label-Free Quantification intensity was used to conduct a t-test and the results were displayed in a volcano plot. This was also done for the unnormalized phosphoproteome. To normalize the phosphosites, the t-test difference of the phospho was subtracted from the t-test difference of the proteome. Sites with a difference of 1 or -1 were considered significantly upregulated or downregulated, respectively. The intensity of the phosphorylation (p)-sites from WT and KO was summed, as depicted in a scatter plot. Regulated p-sites underwent a Fisher’s exact test based on KEGG. Additional graphical visualization was performed in the R environment (version 4.1.1) and GraphPad (version 8.0.1), while figures were edited using Adobe Illustrator. A detailed description can be found in Supplementary methods.

The mass spectrometry proteomics data have been deposited to the ProteomeXchange Consortium via the PRIDE^49^ partner repository with the dataset identifier PXD067740 and 10.6019/PXD067740

### Immunofluorescence

Neurons, plated onto coverslips, were fixed using 4% paraformaldehyde solution, washed and blocked for 1 hour at RT. Primary antibodies were added in 0.1% Triton X-100 and 1% bovine serum albumin overnight at 4°. Finally, the coverslips were washed and mounted to slides using Dako fluorescent mounting medium (S3023, Agilent). Fixed cells were imaged using a standard inverted laser scanning Olympus FV 3000 confocal microscope. Representative images of the cells are shown in the figures with equal and optimal adjustment of brightness and contrast for better visualization.

Primary antibodies include SREBP2 (#28212-1-AP, Proteintech) at 1:500, FLOT1 (#18634, Cell Signaling) at 1:500, DAT (#22524-1-AP, Proteintech) at 1:500 and TOM20 (#sc11415, SantaCruz Biotechnology) at 1:200. Recombinant Clostridium perfringens Perfringolysin O (PFO) tagged with 6xHis (#CSB-EP314820CMB, Hölzel Diagnostika Handels GmbH) and resuspended in 1:1 ratio of water:glycerol to make a 1 mg/mL stock concentration and stored at -20°C. PFO was used at 2.5 μg/mL final concentration onto coverslips. The NR12A dye, a kind gift from Dr. Andrey Klymchenko (University of Strasbourg, France) was used at 40 nM and incubated for 7 minutes. Following the incubation, the media was changed to phenol red free and live-cell imaging was performed using a Leica DMi8 epifluorescence microscope and images were captured using the LASX software. A detailed description can be found in Supplementary methods.

### Immunoblotting

SDS-PAGE gel and protein transfer was followed by blot incubation with primary antibodies – HMGCR (NBP2-66888, Novus Biologicals) at 1:500, SQLE (12544-1-AP, Proteintech) at 1:500, INSIG2 (24766-1-AP, Proteintech) at 1:500, SCAP (PA5-28982, Invitrogen) at 1:500, Miro1 (NBP1-89011, Novus Biologicals) at 1:500, MFN2 (H00009927-M01, Abnova) at 1:500, β3-tubulin (801202, Biolegend) at 1:5000, Vinculin (V9131, Sigma) at 1:2000, GAPDH (CB1001, Sigma) at 1:5000, PKCλ/ι (610208, Biolegend) at 1:500 and PINK1 (846202, Biolegend) at 1:500. Bands were detected with Odyssey CLx (LI-COR) using Image Studio software (LICOR). The band intensities were normalized to a total protein stain – ponceau S solution (#A2935, PanReac AppliChem ITW Reagents) unless otherwise mentioned in the quantification as housekeeping genes – GAPDH and Vinculin were also used for some blots. Image Studio Lite Ver 5.2 (Licor) was used for the quantification of the intensity of bands. A detailed description can be found in Supplementary methods.

### Neurotransmitter Uptake Assay

Neurotransmitter transporter activity in DaNs was measured using the Neurotransmitter Transporter Uptake Assay Kit (#R8174, Molecular Devices) according to the manufacturer’s instructions. Mature DaNs were seeded in Matrigel-coated black, clear bottom 96 well plates prior to the assay at a density of 60,000 cells/well. DaNs were treated with the specific DAT inhibitor GBR 12935 dihydrochloride. Uptake fluorescence was measured using the SpectraMax M2e microplate reader in kinetic mode (Molecular Devices) measuring every 30 s. After the assay, the DaNs were washed and fixed in 4% (v/v) performic acid containing Hoechst to account for cell number in each well.

### Lipidomics

Lipidomics of whole DaNs from independent differentiations (n=3) were performed according to the exact method described in Xicoy et al.^50^

### Mitochondrial Respiration

For the basic mitochondrial stress test, Oxygen Consumption Rate (OCR) and Extracellular Acidification Rate (ECAR)were measured in DaNs using a Seahorse™ XF96 Extracellular Flux Analyzer. Cells were seeded in Matrigel-coated Seahorse cell plates 24-48 h prior to the experiment. A detailed description can be found in Supplementary methods.

### Immunoprecipitation (IP)

A non-denaturing lysis buffer was used containing 20 mM Tris HCl at pH 8, 137 mM NaCl, 10% glycerol, 1% NP-40 and 2 mM EDTA solution. cOmplete protease inhibitor (#11873580001; Sigma) and PhosStop phosphatase inhibitor (#4906837001; Sigma) were added before use. Lysed cells were centrifuged at 15000g for 20 minutes at 4 °C and the supernatant was collected. 1 μg of SCAP polyclonal antibody (PA5-28982, Invitrogen) was added to 1000 μg of total protein and incubated under constant agitation overnight at 4 °C. The next day, 50 μL of Protein A Sepharose beads was added to the solution and incubated under constant agitation for 4 h at 4 °C. A detailed description can be found in Supplementary methods.

### Lipid Order Measurement

NR12A dye, described in Danylchuk et al. ^51^, a kind gift from Dr. Andrey Klymchenko (University of Strasbourg, France) was used at 40 nM and incubated for 7 minutes. Fluorescence was measured using SpectraMax M2e microplate reader with 10 flashes per well in an endpoint reading mode. The excitation wavelength was set at 520 nm and emissions at 560 nm and 630 nm.

### Statistics

For statistical analyses, GraphPad Prism version 8.4.0 was used. The data is presented as mean +/- standard error of mean (SEM) (\**P < 0.05*; \*\**P < 0.005*;

\*\*\**P < 0.0005*; \*\*\*\**P < 0.0001*; nsP *> 0.05*, where ns represents non-significant). Statistical comparisons between more than two conditions were analyzed using an ordinary one-way ANOVA with Tukey’s multiple comparisons tests. Statistical significance between two conditions were determined using unpaired t-test, two-tailed.

## Results

### PINK1 LOF DaNs have reduced SCAP phosphorylation at S822 and S838

To uncover molecular alterations caused by PINK1 LOF, we performed mass spectrometry–based phosphoproteomics in human DaNs carrying either a PINK1 knockout (KO) or the PD patient-derived PINK1 Q456X mutation, each compared to its isogenic control (Fig. 1A-C, Fig. S1A–D). Across all conditions, we consistently quantified around 2000 proteins, with no significant changes (Fig S1A; Supplementary Table 1). A total of 13000 p-sites with a localization probability above 75% were identified, with around 6500 p-sites per condition (Fig S1B; Supplementary Table 1). Among these, SCAP was identified in both PINK1 LOF DaNs having reduced phosphorylation at Ser822 and Ser838 (Supplementary Table 1). Manual inspection of the corresponding annotated MS/MS spectra confirmed the presence and precise localization of these p-sites (Fig S2). Of note, the modified serine residues were assigned as Ser429 and Ser445 in Supplementary Table 1. Mapping of the peptide sequence to UniProt canonical SCAP sequence (Q12770-1) revealed that this residue corresponds to Ser822 and Ser838 respectively having a localization probability > 0.95. We therefore refer to this site as SCAP Ser822 and Ser838 throughout the manuscript. Since no phosphorylation was identified at SCAP Ser 822 and S838 in PINK1 LOF DaNs, it was not possible to include these sites in Fig 1B-C.

**Figure 1:**
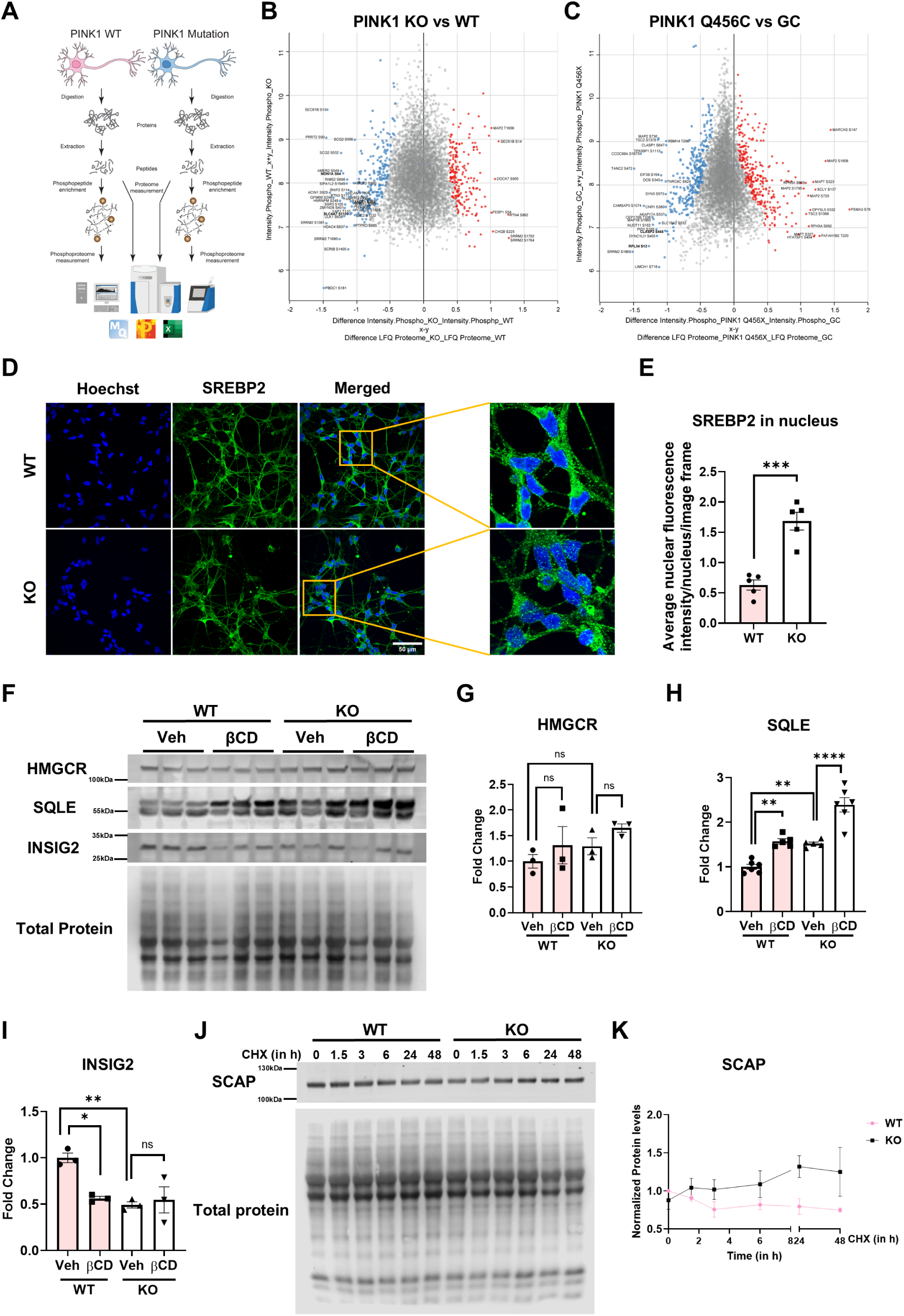
PINK1 LOF dopamine neurons have impaired SCAP phosphorylation at S822 and S838 that increases cholesterol biosynthesis due to increased SCAP stability. A) A schematic workflow for phosphoproteomics. B) Phosphoproteomic differences in PINK1 wildtype (WT) and knockout (KO) dopamine neurons. C) Phosphoproteomic differences in PINK1 Q456X and gene corrected (GC) dopamine neurons. D) Immunofluorescence images showing neuronal localization of SREBP2 (green) in PINK1 WT and KO neurons. E) Quantification of average nuclear fluorescence intensity of SREBP2 in PINK1 WT and KO neurons (n=5). F) Immunoblot showing basal levels of proteins HMGCR, SQLE and INSIG2 in PINK1 WT and ko neurons. G) Quantification of basal HMGCR protein levels (n=3). H) Quantification of basal SQLE protein levels (n=5-6). I) Quantification of basal INSIG2 protein levels (n=3). J) Immunoblot showing SCAP levels in PINK1 WT and KO neurons following a cycloheximide (CHX) treatment to inhibit protein synthesis with timepoint ranging from 0 – 48 hours. K) Quantification of SCAP levels after CHX treatment (n=3). The error bars show the standard error of mean (SEM). For statistical analysis with more than two samples, an ordinary one-way ANOVA was used with Tukey’s multiple comparisons tests. Unpaired t-test (two-tailed) was performed for quantification of the immunofluorescence intensities in (E).

Some of the other differentially phosphorylated proteins identified in PINK1 LOF DaNs included CLASP1 (S598), CLASP2 (S455), MAP1B (S23), and KLC4 (S163), which are associated with microtubule regulation; PAK2 (T169) and DAB2IP (S728), which participate in dendrite development; TBC1D5 (S539) and WDR44 (T94), which function in the endo-lysosomal system; PPME1 (S25) and PPP1R37 (S560), which contribute to protein dephosphorylation and SLC4A7 (S1109) which is involved in the regulation of de novo purine and pyrimidine synthesis downstream of mTORC1 signaling (Supplementary Table 1, Fig 1B-C).

### PINK1 LOF DaNs have increased cholesterol biosynthesis due to increased SCAP stability

Since C-terminal SCAP phosphorylation regulates its stability and interaction with SREBP2^30^, we examined SREBP2 localization. In PINK1 KO DaNs, SREBP2 puncta accumulated in the nucleus compared to the cytosol (Fig. 1D–E), consistent with activation of cholesterol biosynthesis. Immunoblotting revealed upregulation of SQLE and downregulation of INSIG2, whereas HMGCR levels remained unchanged (Fig. 1F–I). As a control we treated DaNs with βCD, a chemical that hydrolyses cholesterol and induces cholesterol biosynthesis.

To show that cholesterol biosynthesis is regulated by PINK1 and not due to clonal artefacts, we expressed WT PINK1 and 3x-kinase-dead (3xKD) mutant PINK1 in PINK1 KO neurons using lentivirus and blotted for HMGCR, SQLE and INSIG2 (Fig S3A-E). Expression of WT PINK1, but not a kinase-dead mutant, normalized SQLE and partially restored INSIG2. However, similar to our observation with βCD treatment, protein levels of HMGCR were unaffected by the expression of WT or 3xKD PINK1.

Reduced phosphorylation in the C-terminus of SCAP in the absence of kinase PKCλ/ι has been previously associated with increased cholesterol biosynthesis due to reduced SCAP degradation^35^. Cycloheximide pulse–chase experiments demonstrated increased SCAP stability in PINK1 KO neurons (Fig. 1J–K), while total SCAP and PKCλ/ι levels were unaffected (Fig. S4A–C). PINK1 did not co-immunoprecipitate with SCAP, suggesting regulation is indirect (Fig. S4D). Together, these results identify impaired SCAP phosphorylation and enhanced cholesterol biosynthesis as downstream consequences of PINK1 loss of function.

## PINK1 LOF increases neuronal cholesterol that is enriched at the plasma membrane and in flotillin-rich lipid rafts

PINK1 has multiple domains including a mitochondrial targeting sequence (MTS), and the TM, the N-terminal (NT), the kinases lobes and the C-terminal region (CTR) (Fig 2A). To gain insights into the pathomechanism, we measured cholesterol content across multiple PINK1 mutations (affecting multiple domains) and LOF models. Free cholesterol was elevated in PINK1 KO and PINK1 Q456X DaNs but not in PINK1 Q126P DaNs (mutation in the N-terminal domain) or in HeLa cells harboring endogenous PINK1 W437X (Fig. 2B–F), suggesting a kinase-domain and neuron-specific phenotype. In PINK1 KO mice, increased striatal cholesterol was observed in young mice but not in the old/adult mice (Fig 2G), consistent with developmental-stage neuronal cholesterol synthesis. In accordance with PD pathology, the ventral midbrain region did not have altered cholesterol both in young and old mice (Fig 2H).

**Figure 2:**
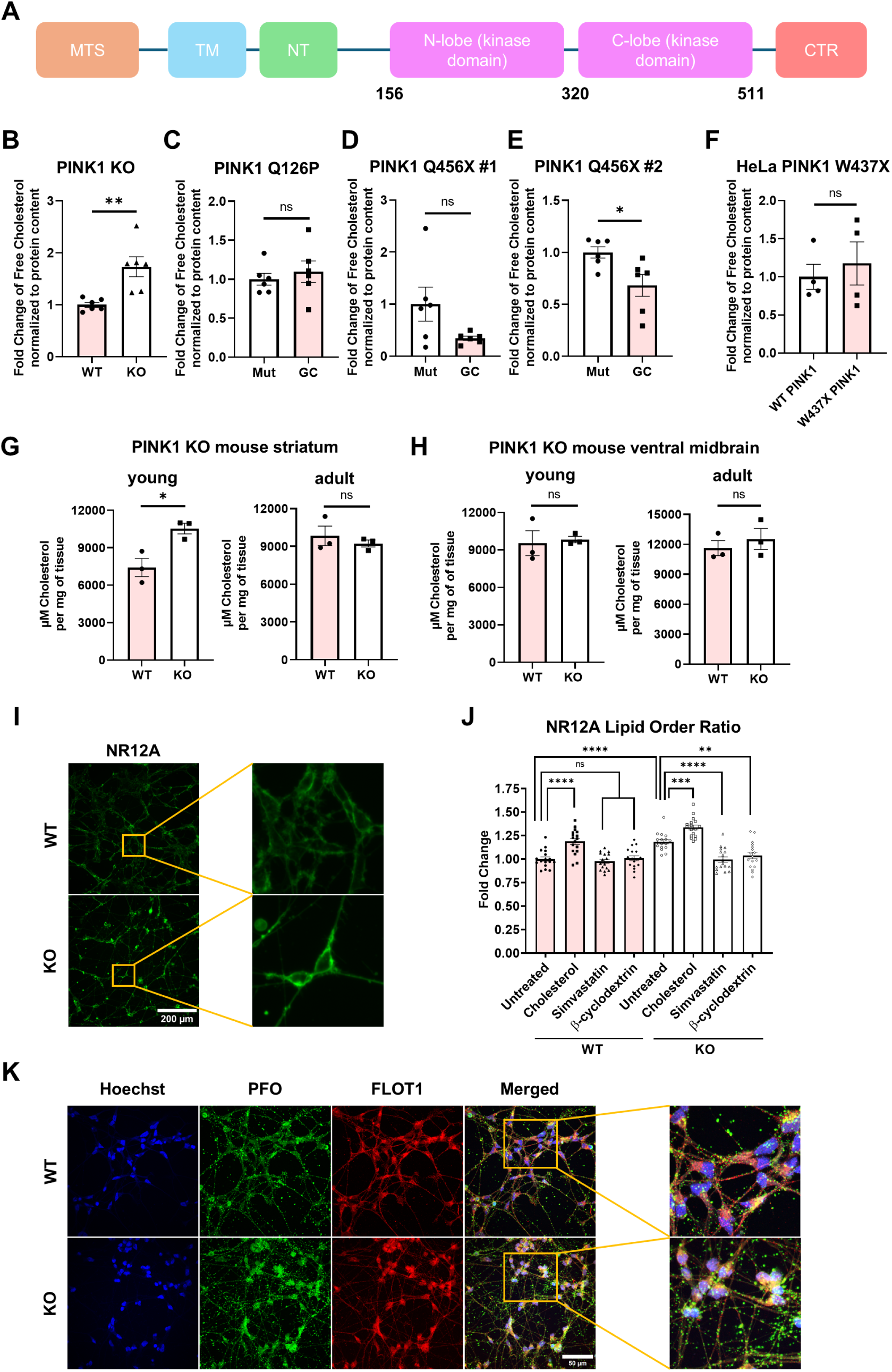
PINK1 LOF human dopamine neurons and young mice striatum exhibit increased free cholesterol which is enriched at the plasma membrane and in FLOT1-rich lipid rafts. A) Schematic for PINK1 domains showing the mitochondrial targeting sequence (MTS) and the kinase domains. Free cholesterol levels of B) PINK1 wildtype (WT) and knockout (KO), C) PINK1 Q126P and gene corrected (GC), D) PINK1 Q456X and GC patient #1, E) PINK1 Q456X and GC patient #2 dopamine neurons (n=6). F) Free cholesterol levels of HeLa cells having endogenous WT PINK1 and PINK1 W437X (n=4). G) Free cholesterol levels of young and old PINK1 WT and KO mice striatum (n=3). H) Free cholesterol levels of young and old PINK1 WT and KO mice ventral midbrain (n=3). I) Live cell epifluorescence images stained with NR12A dye (green) that stains cholesterol at the plasma membranes of PINK1 WT and KO dopamine neurons. J) Plate reader-based quantification of lipid order ratio stained using NR12A dye in live PINK1 WT and KO dopamine neurons (n=17 from 3 independent biological replicates). K) Immunofluorescence images stained with Perfringolysin-O (PFO) in green and Flotillin 1 (FLOT1) in red showing the localization of cholesterol and FLOT1 rich lipid rafts in PINK1 WT and KO dopamine neurons. The error bars show the standard error of mean (SEM). For statistical analysis in (J), an ordinary one-way ANOVA was used with Tukey’s multiple comparisons tests. Unpaired t-test (two-tailed) was performed for quantification of free cholesterol levels in (B-H).

Whole cell lipidomics in PINK1 WT and KO DaNs covering other major lipid classes - phosphatidylglycerol (PG), bis(monoacylglycerol)phosphate (BMP), phosphatidylinositol (PI), phosphatidylethanolamine (PE), phosphatidylserine (PS), phosphatidylcholine (PC), sphingomyelin (SM), lysophosphatidylethanolamine (LPE), cardiolipin (CL), ceramide (Cer) -, revealed no significant alterations (Fig S5).

We next sought to find where in the neurons the level of cholesterol is increased. As the majority of cholesterol is found in the plasma membrane, we used a dye (NR12A) that specifically stains the plasma membrane in live cells. We found that PINK1 KO neurons have increased NR12A binding at the plasma membrane compared to WT neurons (Fig 2I). We also determined the lipid order ratio, using NR12A quantification with a plate reader assay, and found a significant increase in lipid order ratio in PINK1 KO neurons (Fig 2J). Cholesterol was added to the media as a positive control. Additionally, the cholesterol-lowering agents simvastatin and βCD rescued the lipid order ratio.

To visualize if excess cholesterol is also present within the subcellular compartments, we co-stained the neurons with PFO, a toxin that binds to cholesterol^52^, and FLOT1 (a marker for lipid rafts) (Fig 2K). This revealed increased PFO puncta in PINK1 KO neurons that colocalized to FLOT1. Additionally, the signal intensity of FLOT1 is also higher in PINK1 KO neurons suggesting internalization of lipid rafts. By contrast, PFO did not colocalize with mitochondria (Fig. S6), consistent with their normally low cholesterol content. Thus, PINK1 loss specifically drives plasma membrane and FLOT1-rich lipid raft cholesterol accumulation.

### Cholesterol accumulation is independent of mitophagy and does not affect mitochondrial respiration

Since PINK1 regulates mitophagy, we tested whether altered cholesterol levels were linked to this pathway. Antimycin A–induced mitochondrial depolarization failed to trigger degradation of the canonical mitophagy markers Miro1 and MFN2 in PINK1 KO DaNs (Fig. 3A–C), confirming impaired mitophagy.

**Figure 3:**
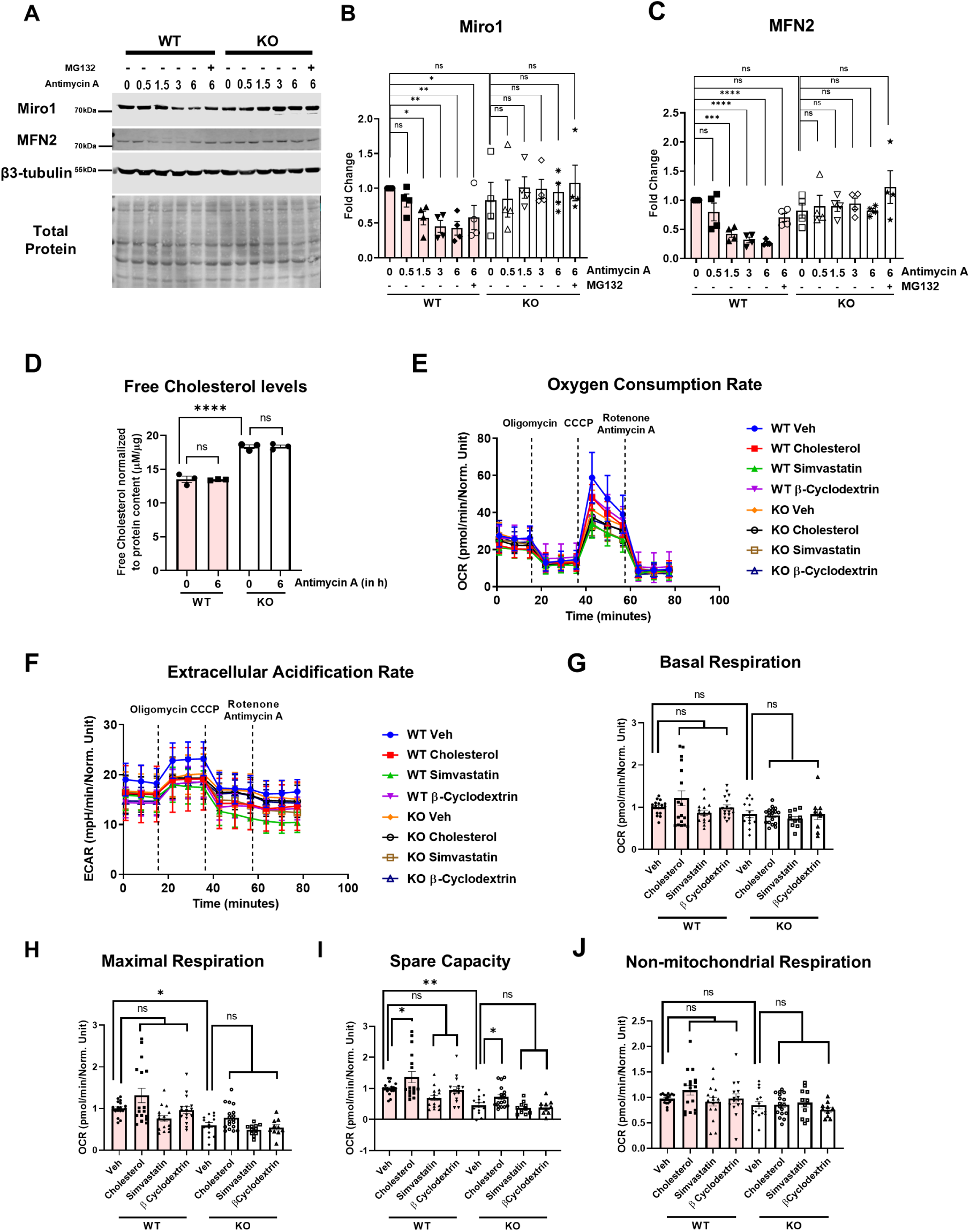
Cholesterol accumulation is independent of mitophagy and does not alter mitochondrial respiration. A) Immunoblot of the mitophagy initiation markers Miro1 and MFN2 in PINK1 wildtype (WT) and knockout (KO) dopamine neurons depolarized using Antimycin A at timepoints ranging 0-6 hours. B) Quantification of Miro1 levels in (A) (n=4). C) Quantification of MFN2 in (A) (n=4). D) Free cholesterol levels measured in PINK1 WT and KO dopamine neurons treated with Antimycin A for 6 hours (n=3). E) Seahorse based mitochondrial stress test to measure Oxygen Consumption Rate (OCR) and F) Extracellular Acidification Rate (ECAR) in PINK WT and KO dopamine neurons treated with cholesterol and cholesterol depleting agents - simvastatin and β-cyclodextrin. G) Quantification of Basal respiration, H) Maximal Respiration, I) Spare capacity and J) Non-mitochondrial respiration in (E) (n=10-18 from 3 independent biological replicates). The error bars show the standard error of mean (SEM). For statistical analysis with more than two samples, an ordinary one-way ANOVA was used with Tukey’s multiple comparisons tests.

The upregulation of cholesterol has been previously shown to inhibit mitophagy^53^ ^54^. However, the levels of cholesterol during mitophagy initiation and whether impaired mitophagy has any effect on cholesterol levels have not been previously studied. Hence, we assessed free cholesterol levels upon mitophagy induction using Antimycin A. However, free cholesterol levels remained unchanged during mitophagy induction (Fig. 3D).

We then assessed whether cholesterol modulates mitochondrial respiration. Seahorse analysis revealed reduced maximal respiration and spare respiratory capacity in PINK1 KO DaNs, while basal and non-mitochondrial respiration were unaffected (Fig. 3E–J). This would suggest that mitochondria are unable to reach their full capacity in the absence of PINK1 as spare respiratory capacity and maximal respiration depends on multiple parameters including the integrity of electron transport chain complexes, the ability of mitochondria to oxidize energetic substrates and mitochondrial health in general^55^. Neither cholesterol depletion nor supplementation altered respiratory parameters. These findings indicate that cholesterol dysregulation in PINK1 neurons occurs independently of mitophagy and does not directly impair basal mitochondrial respiration.

### Excess cholesterol disrupts DAT distribution and neurotransmitter uptake

Cholesterol-rich lipid rafts regulate trafficking of DAT, which contains a cholesterol-binding domain and is internalized through FLOT1-positive microdomains. In PINK1 KO DaNs, DAT redistributed from the plasma membrane to perinuclear regions, partially overlapping with PFO-positive cholesterol puncta (Fig. 4A). Cholesterol depletion with simvastatin or βCD restored normal DAT localization.

**Figure 4:**
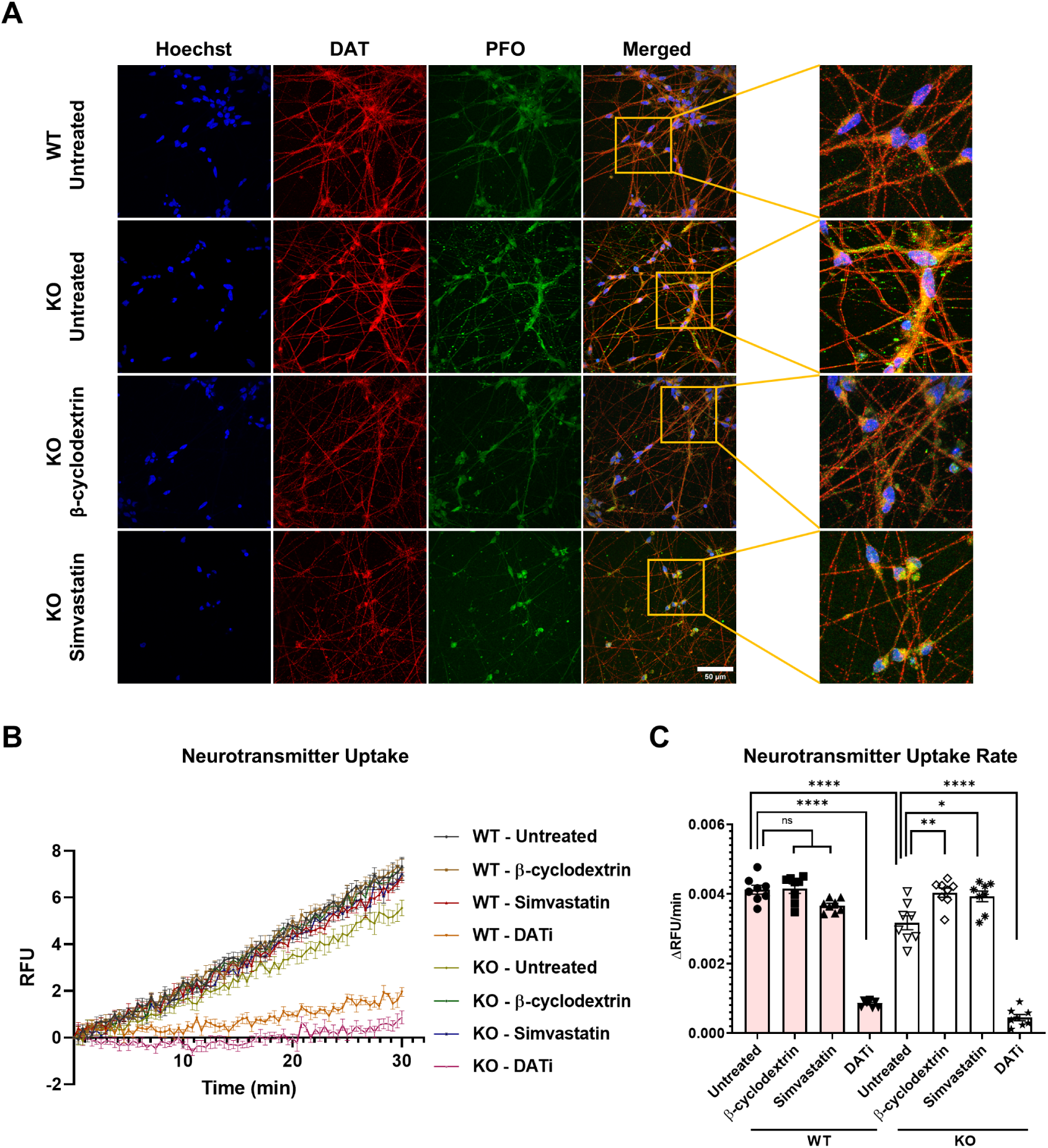
Excess cholesterol disrupts DAT distribution and neurotransmitter uptake. A) Immunofluorescence images showing the distribution of dopamine transporter (DAT) in red and Perfringolysin O (PFO) in green in PINK1 wildtype (WT) and knockout (KO) dopamine neurons either untreated or treated with cholesterol depleting agents - simvastatin and β-cyclodextrin. B) Relative Fluorescence Unit (RFU) of neurotransmitter uptake within a time course of 30 minutes in PINK1 WT and KO dopamine neurons either untreated or treated with simvastatin, β-cyclodextrin or dopamine transporter inhibitor (DATi). C) Quantification of neurotransmitter uptake rate in (B) (n=8 from 4 different neuronal differentiations). The error bars show the standard error of mean (SEM). For statistical analysis with more than two samples, an ordinary one-way ANOVA was used with Tukey’s multiple comparisons tests.

Due to altered DAT distribution and increased cholesterol at the plasma membranes, we performed a neurotransmitter uptake assay (Fig 4B) and found it to be significantly reduced in PINK1 KO DaNs (Fig 4C). This defect was rescued by cholesterol-lowering treatments and was largely DAT dependent, as shown by pharmacological inhibition (Fig 4C). These results demonstrate that cholesterol accumulation in PINK1 LOF neurons disrupts DAT trafficking and impairs dopaminergic function.

## Discussion

By integrating phosphoproteomics and functional assays in multiple PINK1 models, we identify reduced phosphorylation of SCAP at Ser822 and Ser838 as an early event in PINK1-deficient neurons. This reduction stabilizes SCAP, enhances cholesterol biosynthesis, and drives cholesterol accumulation in plasma membrane and FLOT1- rich lipid rafts, with downstream disruption of DAT distribution and neurotransmitter uptake.

SCAP stability is normally controlled by C-terminal phosphorylation, which promotes its ubiquitination and degradation via the proteasome³². We observed reduced phosphorylation of SCAP, increased SREBP2 in the nucleus, and reduced INSIG2 and increased SQLE protein levels in PINK1 LOF neurons, consistent with enhanced cholesterol biosynthesis. Basal SCAP and PKCλ/ι levels were unaffected, raising the possibility that additional kinases or altered post-translational regulation underlie the defect in SCAP phosphorylation.

The physical and biochemical communication between mitochondria and ER at mitochondria-ER contacts sites have been shown to regulate calcium homeostasis^22^, mitochondria dynamics and energetics^56^, apoptosis^57^, and cholesterol metabolism^58^, among others^59^. Identifying the molecular composition and phosphorylation-dependent regulation of these contacts could clarify how PINK1 LOF perturbs cholesterol metabolism in neurons.

A-kinase anchor protein 11 (AKAP11) was another candidate that had reduced phosphorylation at Thr1100 in PINK1 LOF DaNs. It has been reported that astrocytes lacking AKAP11 have upregulated lipid metabolism and an accumulation of esterified cholesterol^60^. While it is not known how phosphorylation of AKAP11 at Thr1100 affects its activity, it has been predicted that PKA and AKAP11 phosphorylates large ribosomal protein 34 (RPL34) at Ser12^61^. Since we also observe a significant downregulation of RPL34 phosphorylation at Ser12 in both PINK1 LOF DaNs, we hypothesize that AKAP11 Thr1100 could impair its activity and may also play a role in cholesterol regulation in PINK1 LOF DaNs.

No differences in cholesterol levels were detected in PINK1 Q126P DaNs. Unlike the Q456X mutation, which resides in the kinase domain, Q126P affects the NT domain of PINK1. This mutation has been reported to disrupt NT–CTR interactions, thereby preventing PINK1 autophosphorylation and stabilization at the outer mitochondrial membrane^62^. Another mutation in the TM region, R98W, alters protein positioning or processing and, in contrast, increases mitophagy through PINK1 accumulation at the outer mitochondrial membrane^63^. Since Q126P does not directly affect the kinase domain, it remains unclear whether its substrate phosphorylation activity is impaired. This underscores the need for further studies to dissect the specific consequences of Q126P and other NT/TM PINK1 mutations.

Cholesterol accumulation has been shown to impair PINK1-Parkin-dependent mitophagy by reducing optineurin recruitment and lysosomal clearance, a phenotype observed in Alzheimer’s disease across several model systems and in *post mortem* brain tissue despite elevated PINK1/Parkin signaling^53^ ^54^. A genome-wide RNAi screen further identified SREBP1 and SREBP2 as key regulators of mitophagy induction, acting by stabilizing PINK1 and promoting Parkin translocation to the mitochondria^64^. These findings suggest that, by disrupting lysosomal clearance, excess cholesterol may impose an additional burden on mitophagy in PINK1 LOF neurons. Moreover, PINK1/Parkin-independent mechanisms for mitochondrial clearance could also be compromised under conditions of cholesterol accumulation.

DAT has six Cholesterol Recognition Amino acid Consensus (CRAC) motifs^65^ through which it can interact with plasma membrane cholesterol/cholesterol-rich lipid rafts. Our study has identified functional consequences of cholesterol accumulation in the plasma membrane. DAT mislocalization to perinuclear regions and its possible partial sequestration into cholesterol-rich rafts impaired neurotransmitter uptake. Restoring the cholesterol balance with simvastatin or βCD corrected these defects, supporting a causal role for cholesterol accumulation. Using super-resolution microscopy, a similar DAT distribution has also been observed in neurons following cholesterol depletion^66^. Interestingly, βCD has also previously been shown to improve differentiation efficiency, bioenergetic profiles and neuronal firing in PINK1 neuronal and midbrain organoid PD models^43^. Our findings align with clinical and preclinical observations that DAT dysregulation precedes neurodegeneration and highlight the contribution of cholesterol homeostasis to dopaminergic vulnerability^67^.

Cholesterol-rich lipid rafts organize signaling and trafficking in neurons^68^. Our observation that FLOT1-positive rafts are increased in PINK1 LOF neurons parallels prior links between lipid rafts and PD-related proteins including α-synuclein, LRRK2, DJ-1, Parkin, and PINK1 itself^69–73^. Notably, loss of Parkin also promotes cholesterol-dependent raft internalization^74^, suggesting convergent lipid regulatory mechanisms across genetic forms of PD.

In summary, we identify reduced SCAP phosphorylation at Ser822 and Ser838 and consequent cholesterol accumulation as early pathogenic features of PINK1 LOF neurons. By linking cholesterol homeostasis to dopaminergic dysfunction, this work expands the biological repertoire of PINK1 beyond mitochondrial quality control and suggests that targeting cholesterol metabolism could modify disease onset or progression in PINK1-related PD.

### Implications and limitations

iPSC-derived DaN model system mimics early stages of neurogenesis, differentiation and maturation, and do not recapitulate aging. We show that these young PINK1 LOF DaNs have increased cholesterol biosynthesis and levels. However, adult neurons do not synthesize their own cholesterol, instead they take in cholesterol predominantly from astrocytes. It is possible that PINK1 astrocytes also exhibit impaired cholesterol homeostasis and future work in astrocytes could be helpful to understand the relevance of cholesterol in PINK1 LOF.

Nonetheless, we observed increased cholesterol levels in the striatum only in the young PINK1 KO mice but not in older mice. This would suggest that this impaired cholesterol homeostasis phenotype could only be present early in the brain when the onset of PD might not be evident. Direct transdifferentiation to human DaNs and/or astrocytes will be useful to understand the relevance of age and cholesterol homeostasis in PINK1 PD.

## Acknowledgements

We thank the study participants, spouses and PD patients who have donated samples to the Hertie Institute for Clinical Brain Research NeuroBiobank. Without their participation, basic research into neuronal mechanisms of disease would not be possible. We thank Nicolas Snaidero, University of Tübingen for providing support with Olympus FV3000 confocal microscopy. We thank the HIH-CIN Imaging Cluster of Microscopy, Core Facility of the Medical Faculty at the University of Tübingen for providing support (especially Olga Oleksiuk) and instrumentation with Leica Dmi8 epifluorescence microscopy. We thank Andrey Klymchenko, University of Strasbourg, France for sharing NR12A dye. We thank Ulrich Rothbauer, University of Tübingen for sharing pLenti6/V5-DEST plasmid with us. pLenti6-DEST PINK1-V5 WT and pLenti6- DEST PINK1-V5 KD were a gift from Mark Cookson, National Institute of Aging, NIH, Bethesda. We thank Jens Schwamborn, Luxembourg Centre for Systems Biomedicine, Luxembourg and Christine Klein, Institute of Neurogenetics, University of Lübeck, Germany for kindly sharing PINK1 Q456X patient and isogenic gene control cell lines with us.

## Author Contributions

Karan Sharma: conceptualization, methodology, validation, formal analysis, investigation, writing—original draft, writing—review and editing, and visualization. Claudia Cavarischia Rega: Methodology, formal analysis, investigation, writing— review and editing, and visualization. Dina Ivanuik – Methodology. Christine Fröhlich (née Bus): Methodology, formal analysis and investigation. Jos F. Brouwers: Methodology and formal analysis. Gerard Martens: Methodology and formal analysis. Philipp Seibler: Resources, writing-review and editing. Javier Jarazo: Resources, writing-review and editing. Thomas Gasser: Resources, writing-review and editing. Daniela M. Vogt Weisenhorn: Resources, writing-review and editing. Boris Macek: Resources, project administration, supervision, writing-review and editing. Julia C. Fitzgerald: conceptualization, methodology, resources, validation, formal analysis, investigation, writing-original draft, writing-review and editing, project administration, supervision, and funding acquisition.

## Supplementary material

This article contains supplemental data.

## Supplementary Figures

**Figure S1:**
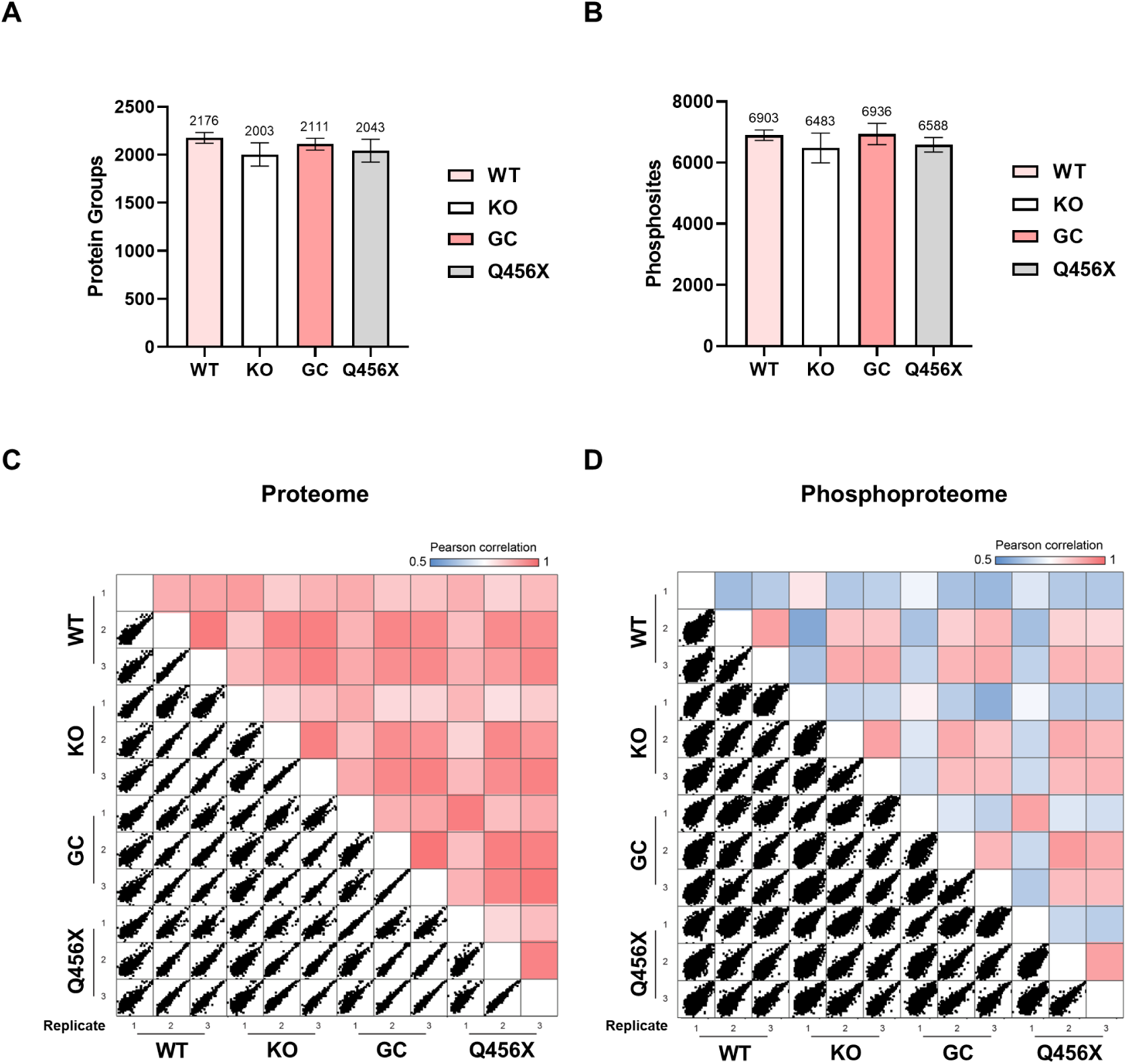
Phosphoproteomics in PINK1 wildtype (WT)/knockout (KO) and PINK1 Q456X/gene corrected (GC) PD patient #2 dopamine neurons. A) Total protein groups and B) phosphosites identified from phosphoproteomic measurements. C) Pearson correlation between PINK1 WT/KO and PINK1 Q456X/GC at the proteome level and D) phosphoproteome level.

**Figure S2:**
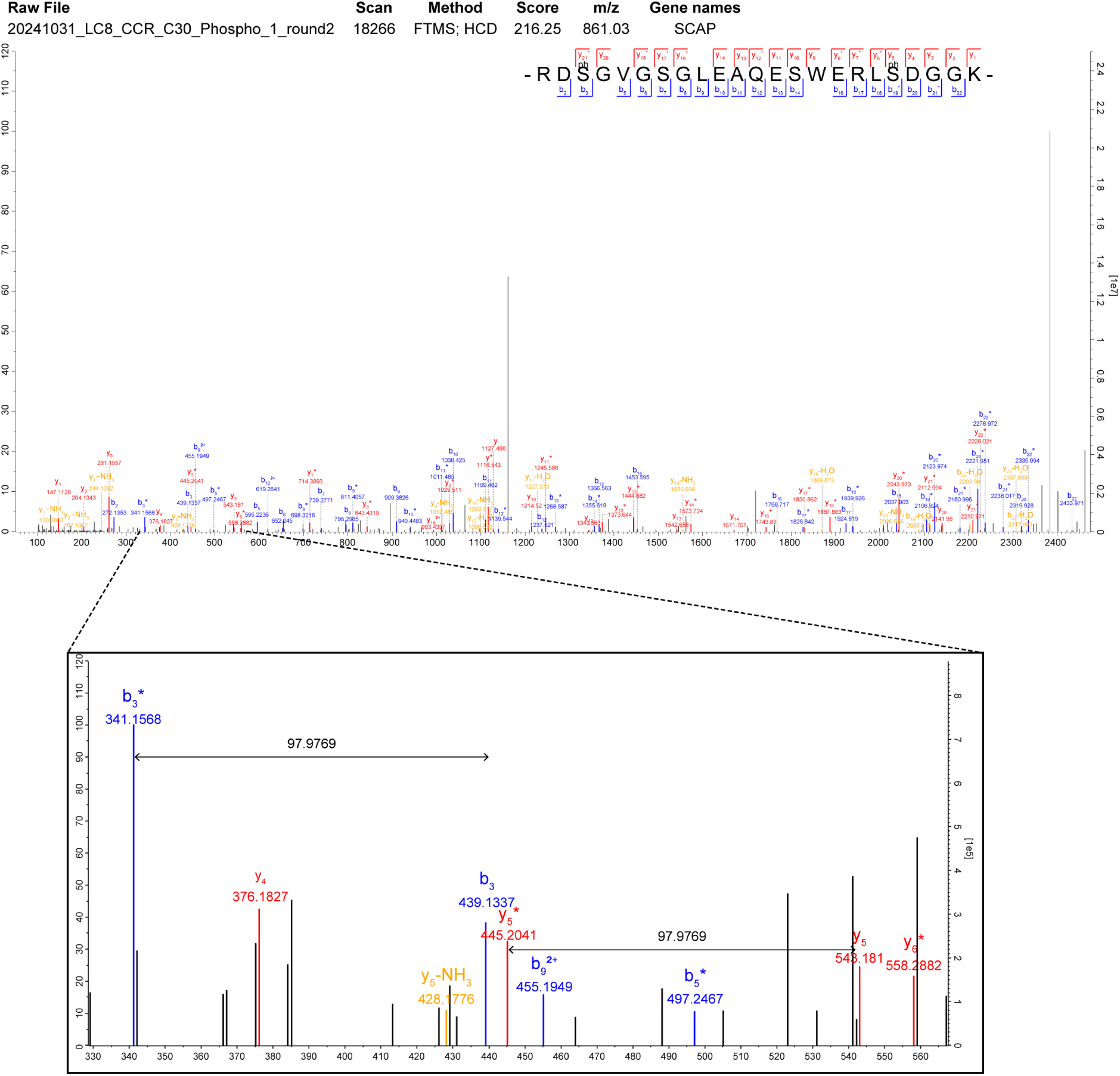
Manual inspection of annotated MS/MS spectra confirming the presence and precise localization of SCAP p-sites.

**Figure S3:**
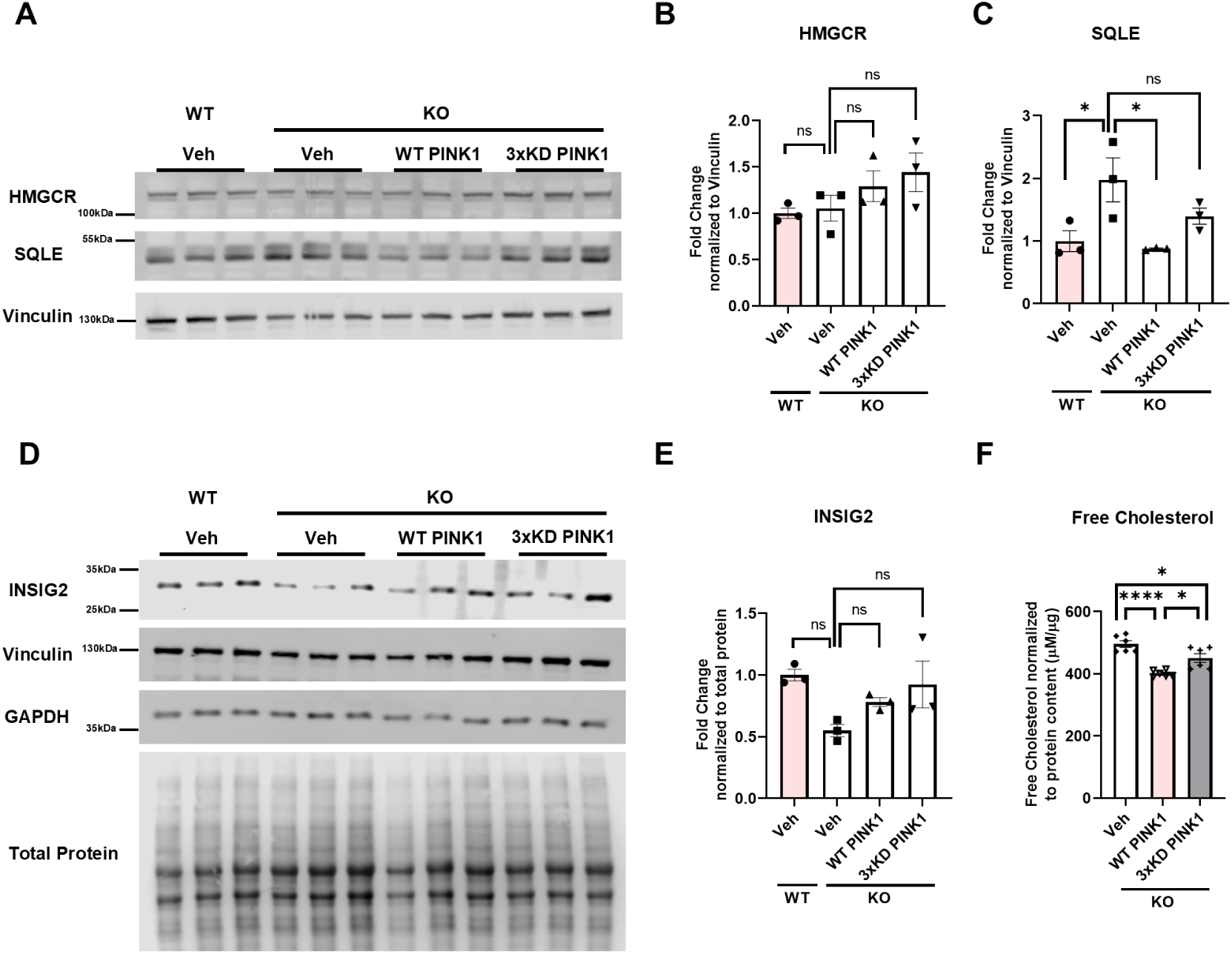
Expression of wildtype (WT) PINK1 virus rescues cholesterol biosynthesis protein levels in PINK1 knockout (KO) dopamine neurons. A) Immunoblot of HMGCR and SQLE in PINK1 WT and KO dopamine neurons treated with either vehicle empty backbone lentivirus (veh), the WT PINK1 lentivirus or the 3x mutant-kinase dead (3xKD) PINK1 lentivirus. B) Quantification of HMGCR in (A) (n=3). B) Quantification of SQLE in (A) (n=3). D) Immunoblot of INSIG2 in PINK1 WT and KO dopamine neurons treated with either vehicle empty backbone lentivirus (veh), the WT PINK1 lentivirus or the 3x mutant-kinase dead (3xKD) PINK1 lentivirus. E) Quantification of INSIG2 in (D) (n=3). F) Free cholesterol measurement in PINK1 KO dopamine neurons treated with either vehicle empty backbone lentivirus (veh), the WT PINK1 lentivirus or the 3x mutant-kinase dead (3xKD) PINK1 lentivirus (n=6). The error bars show the standard error of mean (SEM). For statistical analysis with more than two samples, an ordinary one-way ANOVA was used with Tukey’s multiple comparisons tests.

**Figure S4:**
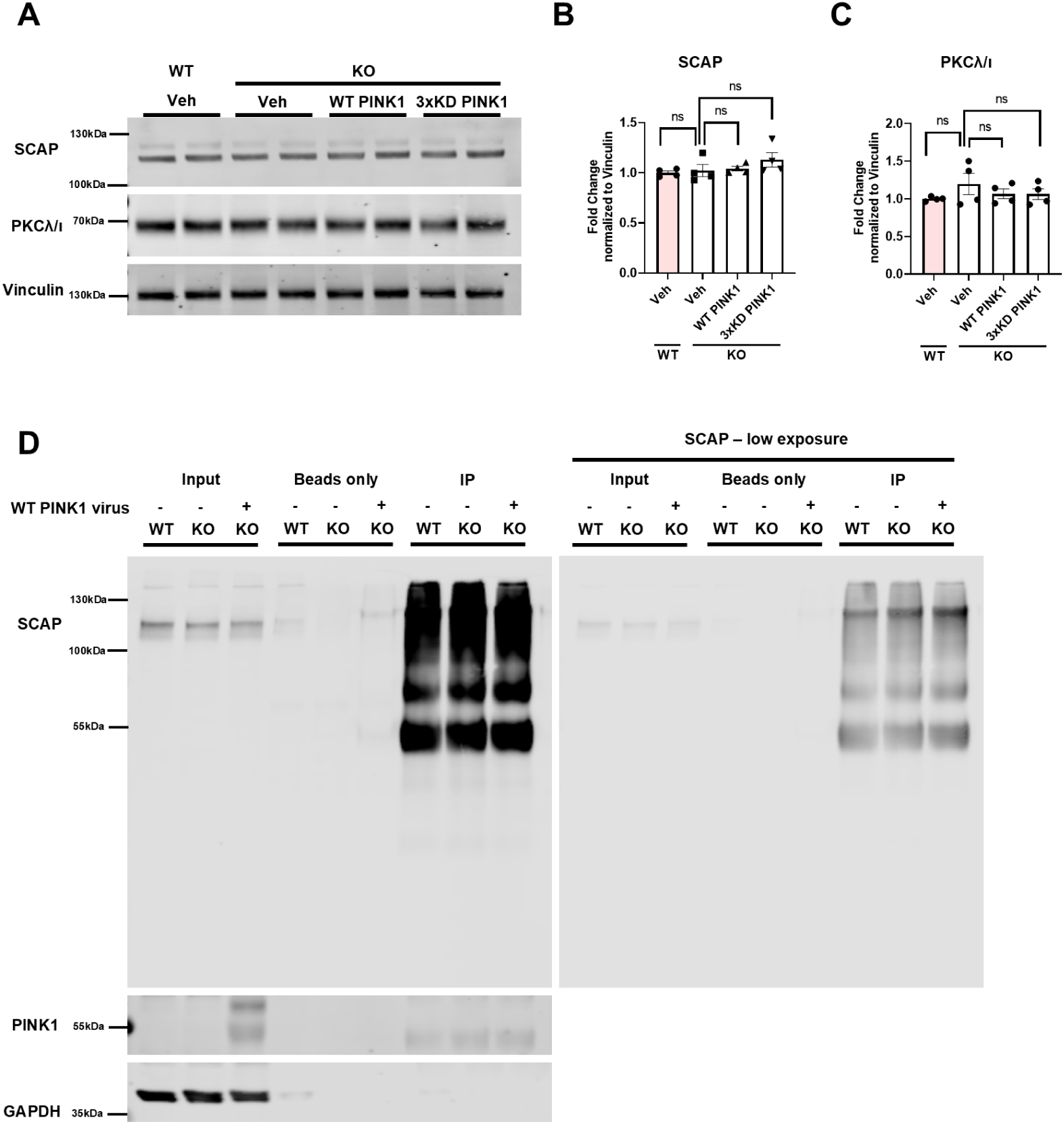
Basal levels of SCAP and PKCλ/ι are unaffected in PINK1 knockout (KO) dopamine neurons and PINK1 does not interact with SCAP. A) Immunoblot of SCAP and PKCλ/ι in PINK1 wildtype (WT) and KO dopamine neurons treated with either vehicle empty backbone lentivirus (veh), the WT PINK1 lentivirus or the 3x mutant-kinase dead (3xKD) PINK1 lentivirus. B) Quantification of SCAP in (A) (n=4). C) Quantification of PKCλ/ι in (A) (n=4). D) Immunoblot of SCAP and PINK1 in PINK1 WT and KO dopamine neurons treated with either vehicle empty backbone lentivirus or the WT PINK1 lentivirus. The blots include the input (whole lysate), immunoprecipitation of beads and immunoprecipitation of SCAP (n=2). The error bars show the standard error of mean (SEM). For statistical analysis with more than two samples, an ordinary one-way ANOVA was used with Tukey’s multiple comparisons tests.

**Figure S5:**
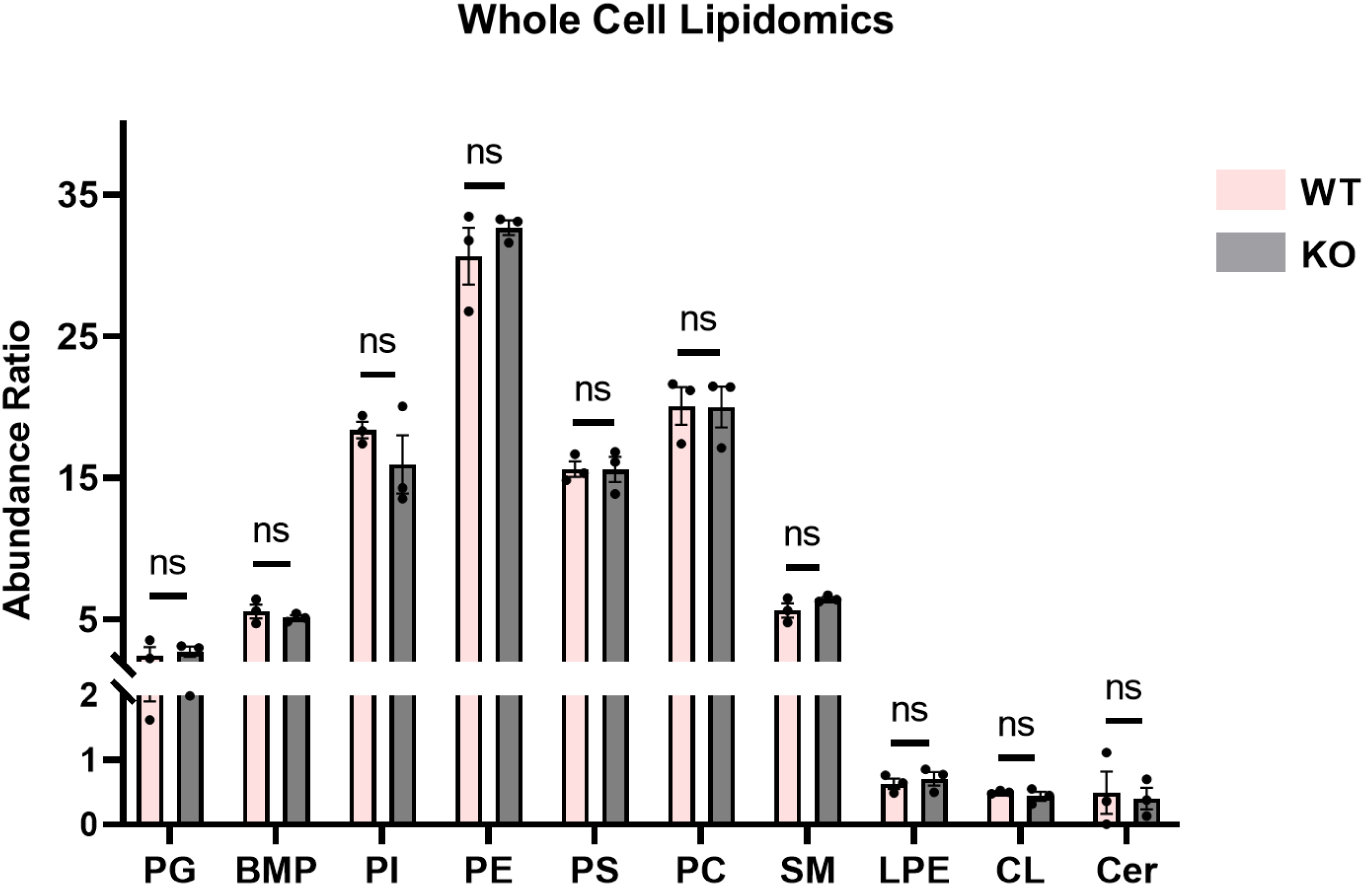
Whole-cell lipidomics in PINK1 wildtype (WT) and knockout (KO) dopamine neurons. Abundance Ratio of major lipid classes – phosphatidylglycerol (PG), bis(monoacylglycerol)phosphate (BMP), phosphatidylinositol (PI), phosphatidylethanolamine (PE), phosphatidylserine (PS), phosphatidylcholine (PC), sphingomyelin (SM), lysophosphatidylethanolamine (LPE), cardiolipin (CL), ceramide (Cer) quantified through whole cell lipidomics in PINK1 WT and KO dopamine neurons (n=3). The error bars show the standard error of mean (SEM). For statistical analysis, an unpaired t-test (two-tailed) was performed.

**Figure S6:**
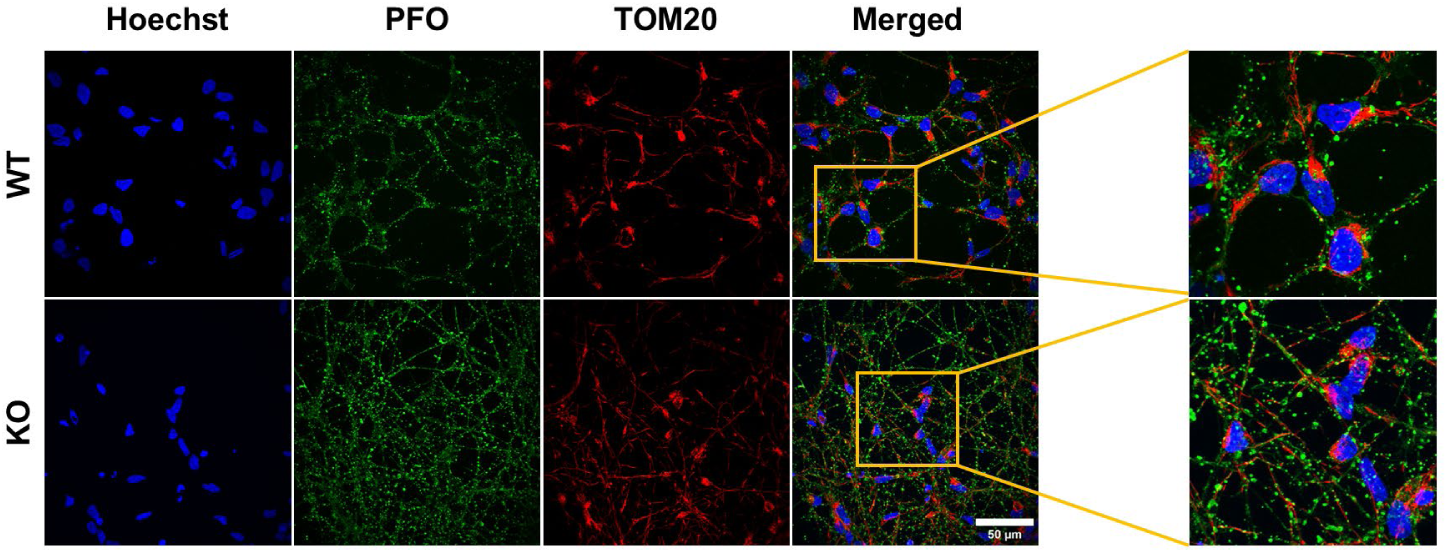
Perfringolysin-O (PFO) stained cholesterol does not colocalize with TOM20 stained mitochondria in dopamine neurons. Immunofluorescence images of cholesterol stained with PFO in green and mitochondria stained with TOM20 in red.

