## Supplementary Methods for "PINK1 regulates cholesterol homeostasis via SCAP phosphorylation in human dopaminergic neurons"

**Generation of PINK1 Q126P Gene Corrected iPSCs**

To generate GC isogenic control line, guide RNA was designed to target exon 1 of the PINK1 gene within the mutation site (GCTCACCTGGATCTCCGGAC). ssODN homology repair construct was designed to be symmetrical to the cleavage site and contained correct sequence (g.155732C>A) and silent mutation in the PAM region (g.155727C>G). Alt-R™ target-specific crRNA oligos, Alt-RHDR ssODN oligos and fluorescent Alt-R™ CRISPR-Cas9 tracrRNA ATTO500 were purchased from IDT Technologies. In order to enhance homologous repair rate IPSCs were treated with non-homologous joining inhibitor Nu7441 (2µM) for 24 hours before and 96 hours after nucleofection. 2 days prior to nucleofection Cas9 expression was induced by treatment with doxycycline (2.5 µg/ml) and continued for 96 hours after nucleofection. iPSCs were dissociated with Accutase™ (Sigma-Aldrich, MO, USA) and gRNA duplex (50 µM) and ssODN (100 µM) were delivered into 10^6^ cells using the Nucleofector Amaxa system (Lonza Biosciences, NC, USA), 3 µl of each. The cells were then re-plated on Matrigel in mTESR1 Plus medium, without antibiotics, supplemented with 10 µM ROCK Inhibitor Y-27632. 24 hours later double positive Cas9-BFP+/sgRNA-ATTO550+ cells were sorted with cell sorter (Sony SH800). Cells were re-plated at low density and subsequently, single colonies were picked up for expansion and transferred onto a 96 well plate. Successful gene correction was assessed by in house Sanger sequencing (PINK1 exon 1 Fw TCGGGCTCGGGCTCCCTAA). The screening of possible off-target effects was performed using CRISPR-Cas9 guide RNA design checker (https://www.idtdna.com/site/order/designtool/index/CRISPR_SEQUENCE). Sequencing primers for off-target effects: (5‘-3‘): NM_001271618.2 PPP1R12C Fw GATCCGCGTCATCAGCAAAC, Rv TGGTCCTGGCTACTGATCCT; NM_145905. HMGA Fw TATGAGAGCTTCCAGGGAGACC, Rv CCCTCATCGAGGGGGTTC; NM_016457.5 PRKD2 Fw TGCTTTTCAAACATGACCCCAC, Rv CAGCACCACCTCCACCAG; NM_015551.2 SUSD5 Fw TCAGACCCGTGAATGCTTCC, Rv TGACAATGGTGGCGATCACA; NG_116786.1 LOC105376345 Fw GAGTGGAATACATGGGTCGC, Rv CAGTTGCCAGTCTCTCTCCAAA; XM_054320707.1 OFD3L2 Fw AGTAGGAGGTGGAGCAGGTT, Rv TGCTTTTCAAACATGACCCCAC; NM_015417.5 SPEF1 Fw ACCCACCGATGGATAGACCA, Rv CTGATTCTGGGGGTGTGGAC.

**Mass-Spectrometry Based Phosphoproteomics**

***Protein Lysis***

Cells were lysed in 1% Triton X-100 in PBS containing cOmplete protease inhibitor (Millipore, 11873580001; Sigma) and PhosStop phosphatase inhibitor (4906837001; Sigma-Aldrich). After lysis, they were centrifuged at 20800 xg for 30 min at 4^o^C. Protein concentration was determined using a micro-BCA kit (#23235 Thermo Scientific).

***Protein precipitation***

Proteins were precipitated with a mix of 100% acetone (ice cold) and 100% MeOH (ice cold) in a ratio of 8:1:1 overnight. The following day, samples were centrifuged at 2000g for 20 minutes at 4°C. The pellets were washed with 80% acetone and then centrifuged again. The pellet was then air-dried and resuspended in DB buffer (6 M urea, 2 M thiourea, and 10 mM Tris pH 8.0). Protein quantification was performed using Bradford reagent, with a standard curve prepared using known concentrations of BSA (Bovine Serum Albumin), and the absorbance was measured at 595 nm. Peptides were purified using solid-phase extraction on Sep-Pak C18 cartridges (Waters, Milford, MA). Phospho-enrichment using titanium dioxide (TiO2) beads.

***Protein in solution digestion***

For each phosphoproteomics experiment, 1 mg per sample was used. Proteins were reduced with 1 mM dithiothreitol (DTT), which was incubated for one hour at room temperature while shaking. To stabilize the reduction, 5.5 mM iodoacetamide (IAA) was added and incubated for one hour at room temperature while shaking in the dark. Samples were pre-digested with LysC (Lysyl Endopeptidase, Wako Chemicals) for 3 hours at RT. Afterwards, four volumes of water were added, and proteins were digested with trypsin (Promega Corporation) overnight. The reaction was stopped the following morning by acidifying the samples to around 0.1% v/v of TFA.

***Phospho enrichment***

Peptides were purified using solid-phase extraction on Sep-Pak C18 cartridges (Waters, Milford, MA). Briefly, the C18 cartridge was activated with 2 mL MeOH and flushed with 2 mL of solvent A* (2% acetonitrile, 1% formic acid) for equilibration. Afterward, the acidified peptide solution was loaded and washed with 2 mL of solvent A (0.1% formic acid). Peptides were eluted with 600 μL of solvent B (80% acetonitrile, 0.5% formic acid). 10 μg per sample was reserved for proteome analysis, while the remainder was used for phospho enrichment. First, titanium dioxide (TiO2) beads were equilibrated in a 1:10 ratio (1 mg of beads per 10 mg of protein) with the loading solution (1 mg beads/100 µL). The peptide solution was incubated with the beads at room temperature for 10 minutes on a spinning carousel. Afterward, the TiO2 beads were spun down, and the supernatant was removed and added to fresh beads prepared in another tube for the next round of enrichment. The TiO2 beads were washed with 100 µL of washing solution I. The supernatant was removed, and the TiO2 beads were resuspended in 100 μL of washing solution II. Then, they were transferred into a C8 tip. The liquid was drained off by centrifugation on a Biofuge Pico at the lowest speed. Samples were then eluted into a fresh 2 mL tube containing 20 µL of 20% TFA using 100 µL of Elution solution I twice and 10 µL of Elution solution II, followed by centrifugation in a Biofuge Pico at the lowest speed. This enrichment process was performed for five consecutive rounds per sample. Eluted samples were subjected to vacuum centrifugation to evaporate ACN.

***Stage tip***

The peptides for proteome and phosphoproteome analysis were desalted and purified using C18 StageTips (Empore). C18 disks were activated with methanol and equilibrated with Solvent A*. The acidified peptides or phosphopeptides were loaded and washed with Solvent A. Peptides were eluted with 50 µl of Solvent B, and ACN was evaporated using vacuum centrifugation. Before loading, the sample volume was adjusted with Solvent A and 10% (v/v) of Solvent A*.

***LC–MS measurement***

Samples were measured on an Exploris 480 mass spectrometer (Thermo Fisher Scientific) online-coupled to a VanquishNeo UHPLC (Thermo Fisher Scientific). Chromatographic separation was performed on a 20 cm long, 75 µm inner diameter analytical HPLC column (ID PicoTip fused silica emitter; New Objective, Berks, UK), packed in-house with ReproSil-Pur C18-AQ 1.9-μm silica beads (Dr Maisch GmbH, Ammerbuch, Germany). Peptides were loaded onto the column at 40°C with a flow rate of 1 µl/min under a maximum backpressure of 1000 bar. For the proteome measurements, peptides were eluted using a 60-minute gradient from 4% to 95% of solvents A (0.1% formic acid) and B (80% acetonitrile in 0.1% formic acid) at a constant flow rate of 200 nl/min.

The mass spectrometer was operated in positive ion mode and data-dependent mode. The acquisition of all full MS scans occurred within a range of 300-1750 m/z at a resolution of 60.000. The twenty most intense multiple-charged ions were selected for HCD fragmentation, with a dynamic exclusion period of 30 seconds. For the proteome measurements, the tandem MS (MS/MS) spectra were acquired at a resolution of 15.000 using a Top20 method. The automatic gain control (AGC) was set to custom, with a normalized AGC target of 50% and an absolute value of 5.000e-4. The maximum injection time was set to auto. For the phosphoproteome measurements, a Top15 method was used, with the resolution of the MS/MS set as 45000. The automatic gain control (AGC) was set to standard. The maximum injection time was set to custom, with a maximum of 220 ms.

***MS data processing***

The raw data were processed using the MaxQuant program (version 2.2.0.0). The raw spectra were searched against the UniProt Homo sapiens database (104556 entries, downloaded 30.01.2024) and commonly observed human contaminants. Cysteine carbamidomethylation was set as a fixed modification, while N-terminal acetylation and oxidation on methionine were selected as variable modifications. The modification Phospho (S/T/Y) was utilized only as a variable modification in the phosphorylation-enriched samples. The peptide mass tolerance was set at 4.5 ppm for MS and 20 ppm for MS/MS. For the tryptic digestion, only two missed cleavages were allowed. The false discovery rate was set to 1% at both the peptide and protein levels. For quantification, both LFQ and iBAQ were selected. All other parameters remained at their default settings.

***Data analysis***

The downstream analysis of MaxQuant output data was performed in Perseus (version 2.0.10.0). Contaminants, reversed hits, and proteins identified only by site were filtered out. Scatter plots were prepared to assess reproducibility between replicates, and Pearson’s correlation was calculated. For the proteome, LFQ intensity was used to conduct a t-test, and the results were displayed in a volcano plot. This was also done for the unnormalized phosphoproteome. To normalize the phosphosites, the t-test difference of the phospho was subtracted from the t-test difference of the proteome. Sites with a difference of 1 or -1 were considered significantly upregulated or downregulated, respectively. The intensity of the p-sites from WT and KO was summed, as depicted in a scatter plot. Regulated p-sites underwent a Fisher's exact test based on KEGG. Additional graphical visualization was performed in the R environment (version 4.1.1) and GraphPad (version 8.0.1), while figures were edited using Adobe Illustrator.

**Immunofluorescence**

Neurons, plated onto coverslips, were fixed using 4% paraformaldehyde solution and washed with PBS. 0.3% Triton X-100 + 1% BSA in PBS was used for permeabilization and blocking and incubated for 1 hour at room temperature. Primary antibodies were added in 0.1% Triton X-100 + 1% BSA overnight at 4^o^. Next day, the cells were washed with 0.1% Tween 20 in PBS. And the secondary antibodies were added to 0.1% Triton X-100 + 1% BSA in PBS for 1 hour at room temperature. Hoechst was then added at 1:10000 (H3569; Molecular Devices, GmbH, Munich, Germany) to stain the nucleus. And then washed with 0.1% Tween 20 in PBS. Finally, the coverslips were washed with PBS and mounted to slides using Dako fluorescent mounting medium (S3023, Agilent). Fixed cells were imaged using a standard inverted laser scanning Olympus FV 3000 confocal microscope. Representative images of the cells are shown in the figures with equal and optimal adjustment of brightness and contrast for better visualization.

Primary antibodies include SREBP2 (#28212-1-AP, Proteintech) at 1:500, FLOT1 (#18634, Cell Signaling) at 1:500, DAT (#22524-1-AP, Proteintech) at 1:500 and TOM20 (#sc11415, SantaCruz Biotechnology) at 1:200. Recombinant Clostridium perfringens Perfringolysin O (PFO) tagged with 6xHis (#CSB-EP314820CMB, Hölzel Diagnostika Handels GmbH) and resuspended in 1:1 ratio of water:glycerol to make a 1mg/mL stock concentration and stored at -20^o^C. PFO was used at 2.5μg/mL final concentration onto coverslips.

Secondary antibodies include Alexa Fluor goat anti-rabbit 647 (#A21245, Invitrogen), donkey anti-mouse 568 (A10037, Invitrogen) and anti-His tag 488 (#652509, Biolegend) at 1:500.

NR12A dye was a kind gift from Andrey Klymchenko, University of Strasbourg, France. It was used at 40nM and incubated for 7 minutes. Following the incubation, the media was changed to phenol red free and live cell imaging was performed using Leica DMi8 epifluorescence microscope and captured using the LASX software.

**Immunoblotting**

Cells were lysed in 1% Triton X-100 + 1% SDS in PBS containing cOmplete protease inhibitor (Millipore, 11873580001; Sigma) and PhosStop phosphatase inhibitor (4906837001; Sigma-Aldrich). After lysis, they were centrifuged at 20800 xg for 30 min at 4^o^C. Protein concentration was determined using a micro-BCA kit (#23235 Thermo Scientific). SDS-PAGE gel and protein transfer was followed by incubation with primary antibodies – HMGCR (NBP2-66888, Novus Biologicals) at 1:500, SQLE (12544-1-AP, Proteintech) at 1:500, INSIG2 (24766-1-AP, Proteintech) at 1:500, SCAP (PA5-28982, Invitrogen) at 1:500, Miro1 (NBP1-89011, Novus Biologicals) at 1:500, MFN2 (H00009927-M01, Abnova) at 1:500, β3-tubulin (801202, Biolegend) at 1:5000, Vinculin (V9131, Sigma) at 1:2000, GAPDH (CB1001, Sigma) at 1:5000, PKCλ/ι (610208, Biolegend) at 1:500 and PINK1 (846202, Biolegend) at 1:500. Secondary antibodies were used at 1:10,000 (#926-32211 and #926-68070; LI-COR Biotechnology - GmbH, Bad Homburg, Germany) and detected with Odyssey CLx (LI-COR) using Image Studio software (LICOR). The band intensities were normalized to a total protein stain – ponceau S solution (#A2935, PanReac AppliChem ITW Reagents) unless otherwise mentioned in the quantification as housekeeping genes – GAPDH and Vinculin were also used for some blots. Image Studio Lite Ver 5.2 (Licor) was used for the quantification of the intensity of bands.

**Mitochondrial Respiration**

For the basic mitochondrial stress test, OCR and ECAR were measured in DaNs using a Seahorse™ XF96 Extracellular Flux Analyzer. Cells were seeded in Matrigel coated Seahorse cell plates 24-48h prior to the experiment. During the experiment cells were treated sequentially with 4μM Oligomycin (#S1478, Selleckchem), 5.6μM CCCP (#C2759, Merck) and 4μM Antimycin A (#A8674-50MG Merck) / 0.8μM Rotenone and the experiment performed according to the manufacturer’s instructions. Following the respiratory analysis, the media was removed, and the cells were fixed with 4% PFA containing Hoechst stain (1:10,000) for 5 minutes. Then washed with PBS. The Hoechst stain is quantified using SpectraMax M2e plate reader. We use Hoechst stain values to normalize the respiration measurements per well.

**Generation of Lentiviral Particles**

A low passage HEK293 FT was cultured in Dulbecco’s Modified Eagles Medium: 4.5 g/L glucose (D6429; Sigma-Aldrich) supplemented with 10% fetal bovine serum (#A5256701, Thermofisher), 1% penicillin/streptomycin (#P0781, Sigma) and 200 μg/mL G418 solution (#G418-B, Capricorn Scientific). These cells were plated onto 10cm dishes. Viral backbone vectors – pMD2.G and psPAX2 and the vectors of interest were pLenti6/V5-DEST, a kind gift from Prof. Ulrich Rothbauer, University of Tübingen. pLenti6-DEST PINK1-V5 WT and pLenti6-DEST PINK1-V5 KD was a gift from Mark Cookson (Addgene plasmid #13319 and #13320 ; <http://n2t.net/addgene:13319>; <http://n2t.net/addgene:13319> ; RRID:Addgene_13319; RRID:Addgene_13320) generated in Beilina et. al.^50^. These vectors were transfected at 3 μg vector of interest, 0.75 μg pMD2.g and 2.25 μg psPAX2 using Roche X-tremeGENE™ HP DNA Transfection Reagent (#6366236001, Merck). The media was collected and concentrated with LentiX concentrator (#631232, Clontech) according to manufacturer’s instructions. Finally, the concentration of the virus thus generated was determined using ZeptoMetrix. Retrotek, HIV-1, p24 antigen ELISA kit (#0801111) following manufacturer’s instructions.

**Immunoprecipitation (IP)**

A non-denaturing lysis buffer was used containing – 20mM Tris HCl at pH 8, 137mM NaCl, 10% glycerol, 1% NP-40 and 2mM EDTA solution. cOmplete protease inhibitor (#11873580001; Sigma) and PhosStop phosphatase inhibitor (#4906837001; Sigma) were added before use. After harvesting cells, the lysis media was added and maintained under constant agitation for 30 minutes at 4^o^C. After which, the lysed cells were centrifuged at 15000g for 20 minutes at 4^o^C and the supernatant was collected. The protein concentration was measured using a micro-BCA kit (#23235 Thermo Scientific). 1 μg of SCAP polyclonal antibody (PA5-28982, Invitrogen) was added to 1000 μg of total protein and incubated under constant agitation overnight at 4^o^C. Next day, 50 μL of Protein A Sepharose beads was added to the solution and incubated under constant agitation for 4 hours at 4^o^C. After the incubation was over, the tubes were centrifuged at 2500g for 2 minutes at 4^o^C and the supernatant was removed. The beads were washed with the lysis buffer and centrifuged under the same conditions and repeated 3 times, removing the supernatant. Finally, 30 μL of 2x NuPAGE™ LDS Sample Buffer (#NP0007, Invitrogen) was added. The sample was heated to 70 ^o^C for 5 minutes and centrifuged again at 5000g for 5 minutes. The supernatant was used as the IP fraction.
